# Conserved effectors underpin the virulence of liverwort-isolated *Pseudomonas* in divergent plants

**DOI:** 10.1101/2024.12.03.626388

**Authors:** Kayla Robinson, Lucia Buric, Kristina Grenz, Khong-Sam Chia, Michelle T. Hulin, Wenbo Ma, Philip Carella

## Abstract

Plant pathogenic *Pseudomonas* species naturally antagonize a diverse range of flowering plants. While emerging research demonstrates that isolates belonging to the *P. syringae* species complex colonize diverse hosts, the extent to which these bacteria naturally infect non-flowering plants like the model liverwort *Marchantia polymorpha* remains unclear. Here, we identify natural associations between *Pseudomonas viridiflava* and the liverwort *Marchantia polymorpha. Pseudomonas* bacteria isolated from diseased liverworts in the wild successfully re-infected *M. polymorpha* in pure culture conditions, producing high *in planta* bacterial densities and causing prominent tissue maceration. Comparative genomic analysis of *Marchantia-*associated *P. viridiflava* identified core virulence machinery like the type-III secretion system (T3SS) and conserved effectors (AvrE and HopM1) that were essential for liverwort infection. Disease assays performed in *Nicotiana benthamiana* further confirmed that liverwort-associated *P. viridiflava* infect flowering plants in an effector-dependent manner. Our work highlights *P. viridiflava* as an effective broad host pathogen that relies on conserved virulence factors to manipulate evolutionarily divergent host plants.

## INTRODUCTION

A diverse range of *Pseudomonas* species interact beneficially, commensally, or detrimentally with plants in natural settings. Amongst this diversity, members of the *Pseudomonas syringae* species complex are prominent pathogens infecting a wide array of flowering plants. In general, the species complex is composed of two distinct clades, a major canonical group of *P. syringae* pathovars infecting crops, and a secondary clade enriched in abiotic isolates but also including related pathogenic species like *P. viridiflava* [1,2]. Experimental interactions between *P. syringae* pathovars and the model flowering plant *Arabidopsis thaliana* have revealed much about the molecular genetic mechanisms underpinning plant immunity and pathogen virulence [3]. However, ecological surveys in North America and Europe demonstrate that *P. viridiflava* is a more prominent pathogen of *Arabidopsis* in natural settings [4,5]. Indeed, *P. viridiflava* has emerged as a successful opportunistic pathogen that infects a wide range of angiosperm hosts [6]. Further dissection of how *P. viridiflava* successfully infects flowering plants like *Arabidopsis* has revealed a key role for pectinolytic enzymes (PELs) that degrade plant cell walls, alongside effector proteins translocated directly into plant cells via the type-III secretion system [7]. Though *P. viridiflava* encode much fewer effectors relative to *P. syringae*, many isolates still harbor the conserved effector locus (CEL) that contains avrE, hopM1, and hopAA1 [1,5].

The vast majority of known interactions between plants and pathogenic *Pseudomonas* species are with hosts belonging to the evolutionarily young angiosperm clade of flowering plants. However, emerging ecological evidence has identified *P. syringae* within aboveground tissues of non-flowering/non-vascular mosses (bryophytes), non-vascular ferns (monilophyte), and even non-plant but photosynthetic lichen [8,9]. In the laboratory, *P. syringae* pv. *tomato* (DC3000) causes chlorotic disease symptoms in the liverwort (bryophyte) *Marchantia polymorpha* [10,11]. Recent research interrogating the virulence profiles of diverse *P. syringae* species complex members further demonstrated that phylogroup 2 isolates enriched in phytotoxins cause severe disease in evolutionarily divergent plants like *Marchantia*, the fern *Ceratopteris richardii*, and the flowering plant *Nicotiana benthamiana* [12]. Intriguingly, species like *P. viridiflava* and *P. cichorii* displayed strong virulence profiles in liverworts and ferns. Whether any of these pathogens naturally associate with liverworts like *M. polymorpha* remains unknown.

Here we report that wild liverworts harbor *P. viridiflava* bacteria that are pathogenic in the model liverwort *M. polymorpha* and the flowering plant *N. benthamiana*. Draft genome sequencing and comparative genomic analyses revealed that infectious isolates encode a canonical T3SS, conserved type-III effector proteins, and the cell wall-degrading pectate lyase (PEL) enzyme. Functional dissection of *P. viridiflava* virulence factors demonstrated the importance of the T3SS and the effectors AvrE and HopM1 for disease establishment in both *Marchantia* and *Nicotiana*. Collectively, our work highlights *P. viridiflava* as an ecologically relevant pathogen of *Marchantia* liverworts and demonstrates that conserved effectors are critical for the manipulation of distantly related host plants.

## RESULTS

### Wild *Marchantia polymorpha* harbor plant pathogenic *Pseudomonas*

To determine whether liverworts naturally harbor plant pathogenic *Pseudomonas*, we first isolated endophytic bacteria from the aerial gametangiophores (sexual reproductive structures) of wild *Marchantia* liverworts exhibiting disease symptoms like tissue pigmentation, browning, and maceration (Fig. 1AB). To enrich for pathogenic *Pseudomonads*, endophytes were selected on KB media supplemented with the antibiotic nitrofurantoin [5]. Next, we screened 40 *Marchantia* endophytes (Fig. 1C), which we named MPG (“*Marchantia, purple gametophore”)*, for their ability to re-infect wildtype Tak-1 liverworts in axenic conditions (Fig. S1). This revealed a number of isolates that produced disease symptoms ranging from tissue darkening and maceration to chlorosis (yellowing) in the vegetative plant body (thallus). We further interrogated bacterial growth and disease symptom development using a subset of disease-promoting MPG isolates alongside a non-infectious strain identified from our initial screen in *Marchantia* (MPG01). As expected, MPG01 failed to produce disease symptoms and grew to lower *in planta* bacterial densities compared to virulent MPG strains (Fig. 1D). Collectively, this demonstrates that *Pseudomonas* bacteria naturally infect *M. polymorpha* liverworts.

**Figure 1.**
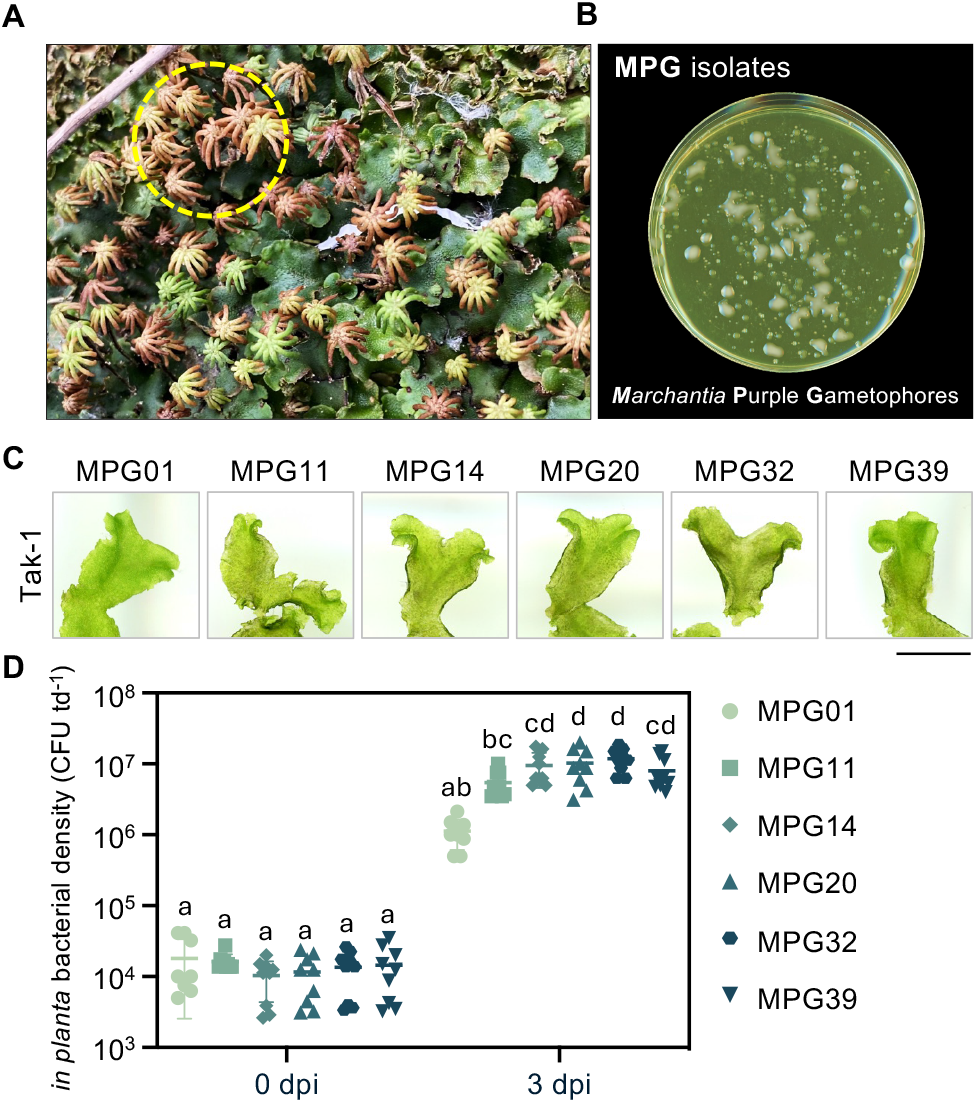
Wild *Marchantia* harbor plant pathogenic *Pseudomonas* sp. (A) Wild *Marchantia* liverworts showing disease-like symptoms in their aerial organs. Endophytic bacteria were isolated from symptomatic archegoniophore heads (dashed circle). (B) Example of wild *Pseudomonas* MPG (Marchantia, purple gametophore) isolates growing KB media supplemented with the *Pseudomonas-*selective antibiotic nitrofurantoin. (C) Symptom development of axenically grown wildtype *Marchantia polymorpha* (Tak-1) infected with MPG isolates. Symptoms were recorded 7 days post infection (dpi) (n ≥ 8). Performed at least three times with similar results. Scale bars = 1 cm. (D) *In planta* bacterial densities of MPG isolates in *M. polymorpha* (Tak-1) at 0 and 3 dpi. Each datapoint represents a single biological replicate (n ≥ 8), with three biological replicates included for each of three experimental replicates (total of 9 datapoints). Different letters signify statistically significant differences in bacterial densities (ANOVA, Tukey’s HSD, p < 0.05). Error bars represent standard deviation.

### Liverwort-isolated *Pseudomonas viridiflava* belong to the *P. syringae* species complex

Next, we generated draft genomes to delineate species relationships between *Marchantia* endophytes and to assess their virulence factor repertoires (Figure S2). A maximum likelihood species phylogeny of 706 concatenated orthologs shared between each MPG isolate and diverse representatives from across the *P. syringae* complex revealed that virulent MPG isolates were situated within phylogroup (PG) 7 alongside *P. viridiflava*, whereas non-infectious MPG01 was excluded from the complex similar to a *P. putida* outgroup (Fig. 2A). Taking a sequence-homology based approach using existing knowledge from *P. syringae* and *P. viridiflava*, we then resolved the virulence factor repertoires of infectious MPG isolates. In each *Marchantia-*derived isolate, we identified the cell wall degrading pectate lyase enzyme PEL alongside the type-III effectors AvrE, HopM1, HopAA1, HopA1, and HopB2. Phylogenetic analysis of each virulence factor highlighted similarities with PG7 isolates like the *P. viridiflava* type strain ICMP2848 or the closely-related *P. syringae* pv. *ribicola* and *P. syringae* pv. *primulae* (Figures S3-S8), which are potentially misidentified as *P. syringae* rather than *P. viridiflava* [13]. Moreover, the presence of multiple CEL effectors indicates that these isolates harbor the more canonical tripartite pathogenicity island (T-PAI) rather than the reduced single pathogenicity island (S-PAI) often associated with *P. viridiflava* [6]. Collectively, the data suggest that PG7 *P. viridiflava* are natural antagonists of *Marchantia* that harbor conserved virulence factors.

**Figure 2.**
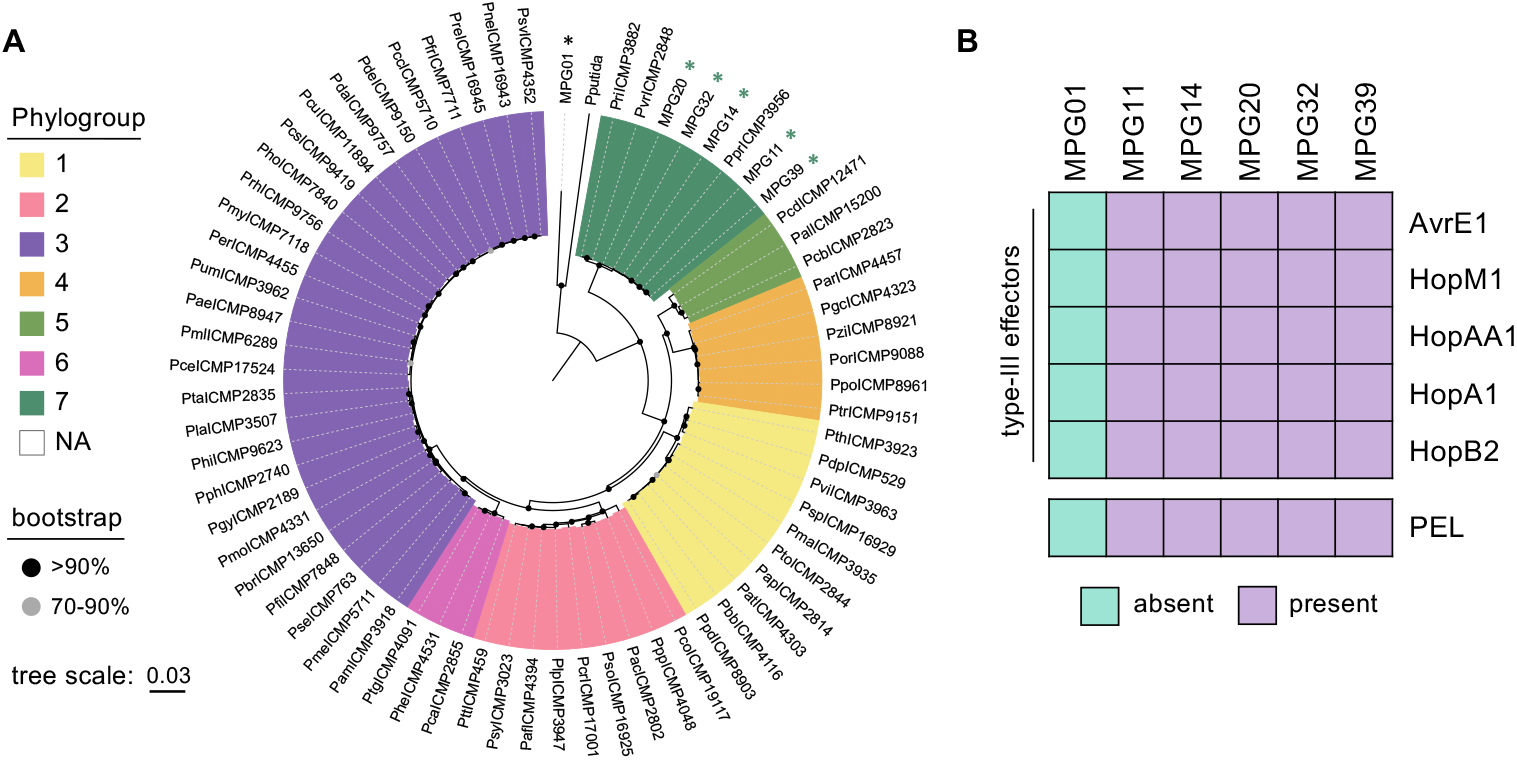
Liverwort-associated *Pseudomonas* belong to phylogroup 7 of the *P. syringae* species complex. (A) Maximum likelihood phylogeny of concatenated single-copy orthologs shared between MPG isolates, diverse *P. syringae* species complex members, and a *P. putida* outgroup (genomes listed in Table S1). Different colors represent different phylogroups of the species complex. Tree scale = substitutions/site. (B) Presence or absence of type-III effectors or pectate lyase (PEL) virulence factors in liverwort-associated MPG isolate genomes.

### Liverwort infection relies on the T3SS and Effectors, not PEL

To determine the functional relevance of *P. viridiflava* virulence factors for liverwort infection we generated and characterized knockouts of the *hrcC* gene essential for T3SS function (effector delivery) or the pectinolytic enzyme *pel* in the *P. viridiflava* MPG32 background (Figure S9). Infection assays in Tak-1 liverworts demonstrated that ΔhrcC mutants failed to promote disease symptoms or appreciably proliferate *in planta*, whereas Δpel mutants were indistinguishable from wildtype (Figure 3AB). Importantly, Δpel bacteria were unable to cause pectinolytic-dependent rotting in potato tubers, resembling the uninoculated mock control treatment (Figure 3C). By comparison, both MPG32^WT^ and ΔhrcC caused severe rotting, confirming that *P. viridiflava* MPG32 relies on PEL for pectinolytic activity. Together, the data suggest that T3SS-delivered effectors rather than PEL activity is central to *P. viridiflava* virulence in liverworts.

**Figure 3.**
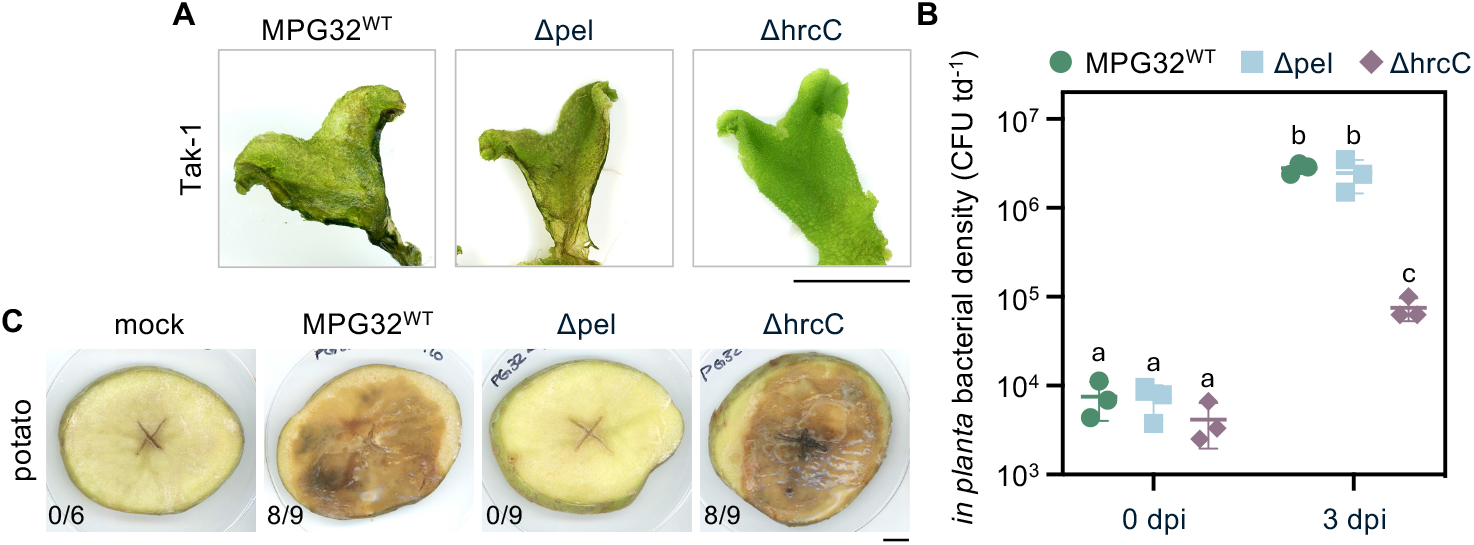
The type-III secretion system is essential for MPG virulence in *Marchantia*. (A) Disease symptom development in wildtype *Marchantia polymorpha* (Tak-1) infected with MPG32^WT^, Δpel, and ΔhrcC mutants. Symptoms were recorded 7 days post infection (dpi) (n ≥ 8). Performed at least three times with similar results. Scale bars = 1 cm. (B) *In planta* bacterial densities of MPG32^WT^, Δpel, and ΔhrcC mutants. The plot depicts a single experimental replicate where each datapoint represents a biological replicate (n ≥ 8). Different letters signify statistically significant differences in bacterial densities (Multiple pairwise comparison, Student’s T-tests, p < 0.05). Error bars represent standard deviation. Performed three times with similar results. (C) Pectate lyase activity tests of MPG32^WT^, Δpel, ΔhrcC mutants relative to a mock untreated control. Potato slices were imaged 3 days post inoculation. Numbers indicate the potato slices exhibiting pectate lyase-dependent maceration relative to the total number of inoculated samples per treatment. Performed twice with similar results. Scale bars = 1 cm.

### Conserved AvrE and HopM1 effectors are essential for infection in *Marchantia*

The importance of the T3SS for *P. viridiflava* virulence in *Marchantia* suggests that type-III effectors are crucial for disease establishment. Amongst the relatively small effector repertoire of MPG isolates, AvrE and HopM1 stand out as conserved and broadly distributed effectors that are generally essential for *P. syringae* infection in angiosperms [14–16]. Based on AlphaFold3 protein structure modelling, both AvrE and HopM1 from *P. viridiflava* MPG32 share considerable structural homology to corresponding effector alleles from well characterized strains like *P. syringae* pv. *tomato* DC3000 and *P. syringae* pv. *syringae* B728a (Figure 4A). To evaluate their contribution to virulence in *Marchantia*, we generated and performed infection assays with a double ΔavrE/hopM1 knockout for comparison against a wild-type *P. viridiflava* MPG32^WT^ control (Figure S9). Consistent with their importance in flowering plants, the *P. viridiflava* ΔavrE/hopM1 double mutant was significantly less virulent in *M. polymorpha* Tak-1 compared to MPG32^WT^. This further underscores the importance of the conserved AvrE and HopM1 effectors for disease establishment across distantly related host plants.

**Figure 4.**
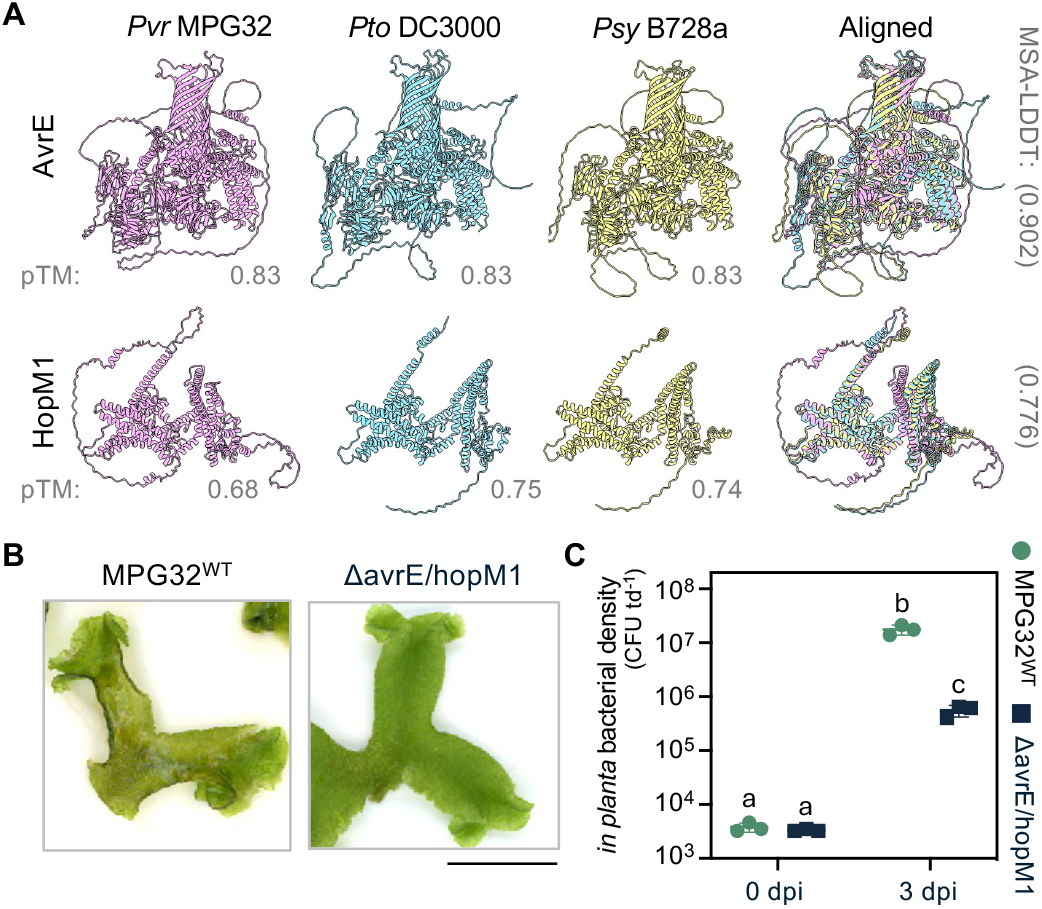
The conserved AvrE and HopM1 effectors are critical for MPG virulence in Marchantia. (A) Structure model similarity of *Pseudomonas viridiflava* MPG32 AvrE and HopM1 effectors against corresponding alleles in *P. syringae* pv. *tomato* DC3000 and *P. syringae* pv. *syringae* B728a. AlphaFold3 generated models are shown alongside pTM confidence scores for each model. FoldMason-generated structural similarity/alignment scores (MSA-LDDT) are provided next to merged protein models. (B) Disease symptom development of wildtype *Marchantia polymorpha* (Tak-1) infected with MPG32^WT^ or the ΔavrE/hopM1 double mutant. Symptoms were recorded 7 days post infection (dpi) (n ≥ 8). Performed at least three times with similar results. Scale bars = 1 cm. (C) *In planta* bacterial densities of MPG32^WT^ or ΔavrE/hopM1 double mutants in *M. polymorpha* (Tak-1) at 0 and 3 dpi. The plot depicts a single experimental replicate where each datapoint represents a biological replicate (n ≥ 8). Different letters signify statistically significant differences in bacterial densities (Multiple pairwise comparison, Student’s T-tests, p < 0.05). Error bars represent standard deviation. Performed three times with similar results.

### Liverwort-isolated *P. viridiflava* infect *Nicotiana benthamiana*

Virulent *P. viridiflava* have been isolated from a diverse range of flowering plants [6]. To test whether *Marchantia*-associated *P. viridiflava* exhibit broad host virulence in a flowering plant, we assessed the virulence profiles of the 6 genome-sequenced MPG isolates in *Nicotiana benthamiana* leaves. Similar to liverworts, all isolates with exception to MPG01 exhibited strong disease symptoms and grew to high *in planta* bacterial levels (Figure 5A). Further analysis of *P. viridiflava* MPG32 mutants in *N. benthamiana* again demonstrated key roles in virulence for the T3SS and AvrE/HopM1 but not PEL (Figure 5BC). Collectively, the data demonstrate that liverwort-associated *P. viridiflava* rely on conserved type-III effectors to cause disease in nonflowering and flowering plants alike.

**Figure 5.**
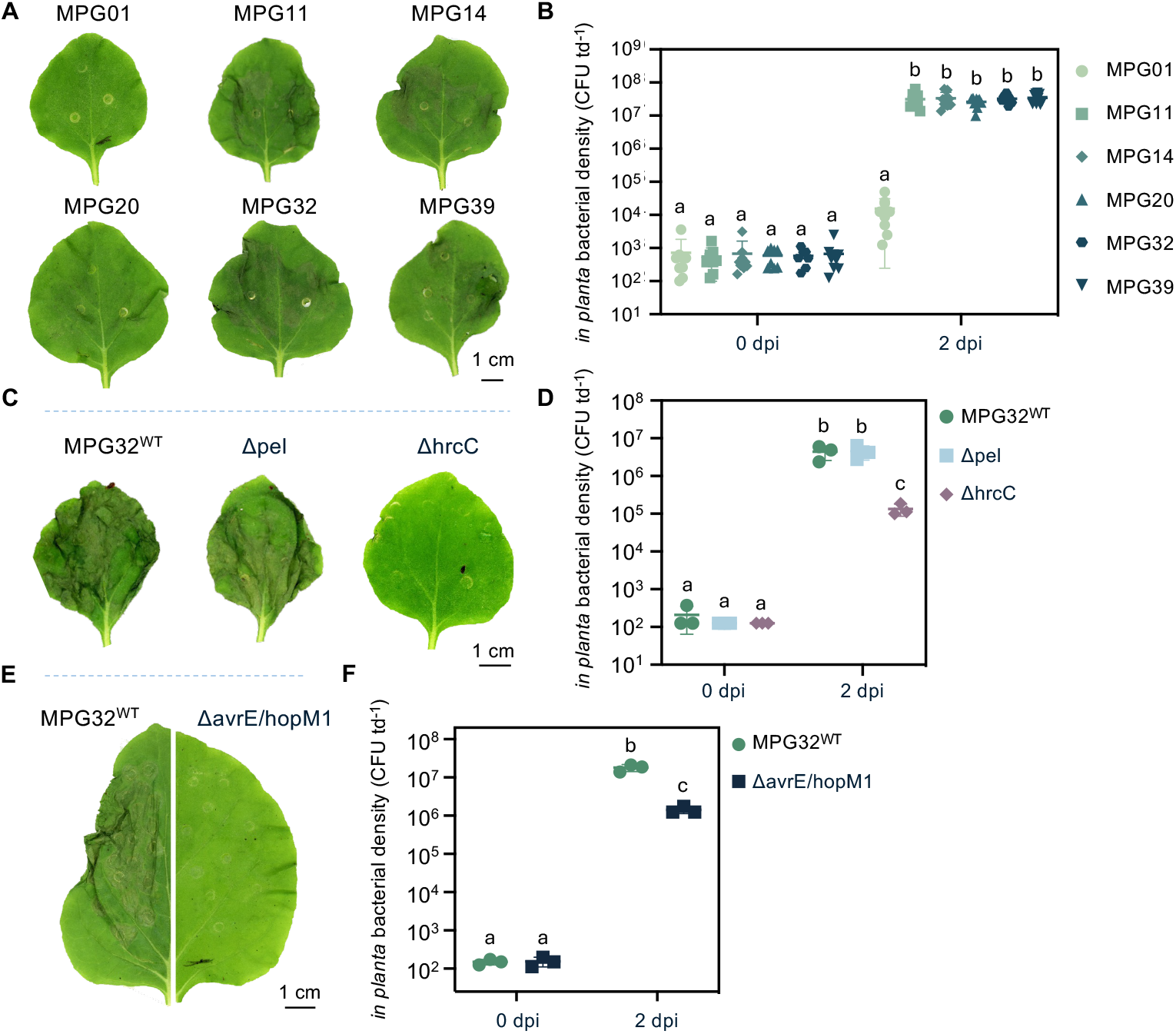
Liverwort-associated MPG infect the flowering plant *Nicotiana benthamiana*. (A) Disease symptom development of wild-type *Nicotiana benthamiana* infected with MPG isolates at 2 dpi. Performed three times with similar results. Scale bars = 1 cm. (B) *In planta* bacterial densities of MPG isolates in *N. benthamiana* leaves at 0 and 2 dpi. Each datapoint represents a single biological replicate (n ≥ 8), with three biological replicates included for each of three experimental replicates (total of 9 datapoints). Different letters signify statistically significant differences in bacterial densities (ANOVA, Tukey’s HSD, p < 0.05). Error bars represent standard deviation. (C) Disease symptom development of *N. benthamiana* infected with MPG32^WT^, Δpel, and ΔhrcC mutants at 2 dpi. Performed three times with similar results. Scale bars = 1 cm. (D) *In planta* bacterial densities of MPG32, Δpel, and ΔhrcC in *N. benthamiana* leaves at 0 and 2 dpi. The plot depicts a single experimental replicate where each datapoint represents a biological replicate (n ≥ 8). Different letters signify statistically significant differences in bacterial densities (Multiple pairwise comparison, Student’s T-tests, p < 0.05). Error bars represent standard deviation. Performed three times with similar results. (E) Disease symptom development of MPG32^WT^ and the ΔavrE/hopM1 double mutant in *N. benthamiana* leaves at 2 dpi. Performed three times with similar results. Scale bars = 1 cm. (F) *In planta* bacterial densities of MPG32 and the ΔavrE/hopM1 double mutant in *N. benthamiana* leaves at 0 and 2 dpi. The plot depicts a single experimental replicate where each datapoint represents a biological replicate (n ≥ 8). Different letters signify statistically significant differences in bacterial densities (Multiple pairwise comparison, Student’s T-tests, p < 0.05). Error bars represent standard deviation. Performed three times with similar results.

## DISCUSSION

In this study, we demonstrate that *P. viridiflava* is an effective pathogen of the liverwort *M. polymorpha*. Importantly, endophytic bacteria isolated from wild liverworts caused disease in naïve lab-grown *Marchantia*, thus fulfilling Koch’s postulates for microbial pathogenesis [17]. Previous microbiome studies of wild *M. polymorpha* and the related species *M. paleacea* identified a range of bacteria likely involved in plant growth promotion [18]. By comparison, investigations of *M. polymorpha* fungal endophytes revealed both beneficial and detrimental isolates, of which *Xylaria* and *Colletotrichum* species were the most pathogenic in lab-based infection assays [19]. The fungal pathogen *Irpex lacteus* has also been identified from wild *Marchantia*, which likely adopts a necrotrophic (tissue-killing) lifestyle [20]. Collectively, these works begin to resolve the diversity of pathogenic microbes that naturally antagonize non-flowering liverworts.

Liverwort-associated *P. viridiflava* retain the capacity to infect the flowering plant *N. benthamiana*, which highlights the broad host nature of this pathogen. This brings much needed ecological context to previous studies demonstrating strong virulence of angiosperm-derived *P. syringae, P. cichorii*, and *P. viridiflava* in *Marchantia* [10–12]. Indeed, many pathogens and pests originally associated with flowering plants are capable of infecting liverworts, including the oomycete *Phytophthora palmivora* [21], the fungus *Fusarium oxysporum* [22], *Tobacco Mosaic Virus* (TMV) [23], and insects like *Armadillidium vulgare* [24,25] and *Scatopsciara cunicularius* [26]. Similar to liverworts, mosses also succumb to angiosperm-isolated pathogens [27–29] and naturally harbor diverse microbes from genera known to impact crops [8,30,31], including a strain of *Fusarium avenaceum* that caused necrotic disease when re-introduced to the moss *P. patens* or the flowering plants barley, tomato, and carrot [32]. These studies demonstrate that nonflowering plants are relevant model systems to understand plant-pathogen interactions, and hint toward the potential for bryophytes to act as natural reservoirs for angiosperm pathogens.

Functional dissection of *P. viridiflava* virulence in *Marchantia* revealed a strong reliance on T3SS-delivered effectors rather than the pectinolytic cell wall degrading enzyme PEL. Previous research demonstrated that PEL enzymatic activity was required for tissue maceration (soft-rotting) and disease establishment in Arabidopsis and pepper [7,33]. Intriguingly, PEL activity is higher in *P. viridiflava* isolates containing the single pathogenicity island (S-PAI) and a single type-III effector (AvrE) compared to those encoding the canonical tripartite pathogenicity island (T-PAI) and multiple effectors [7]. Liverwort *P. viridiflava* harbor multiple effectors that are generally absent from S-PAI isolates [5,7,34], which may explain their reliance on the T3SS rather than PEL. By contrast, the conserved effectors AvrE and HopM1 were essential for *P. viridiflava* virulence. In flowering plants, both AvrE and HopM1 promote water soaking (hydration) in the host apoplast to support bacterial proliferation and suppress plant immunity. This is achieved by interfering with host proteins regulating immunity and water homeostasis [15,16,35]. Recent cell physiology and structural biochemistry studies also provide evidence that AvrE acts as a water and solute permeable channel directly in plant membranes [36]. Given the importance of AvrE and HopM1 for *P. viridiflava* infection in *Marchantia* and *Nicotiana*, it is likely that these effectors retain their infection-promoting molecular functions across divergent plant hosts. Future research identifying conserved susceptibility hubs perturbed by AvrE, HopM1, and even other type-III effectors will reveal macroevolutionary underpinnings of *Pseudomonas* virulence in land plants. Whether additional small molecule toxins or other virulence factors also play roles in *P. viridiflava* virulence remains to be investigated.

Our work highlights *P. viridiflava* as a natural pathogen of liverworts that successfully antagonizes *M. polymorpha* and the flowering plant *N. benthamiana*. While this study focused on virulence factors enabling pathogenicity in divergent hosts, future research leveraging plant diversity may identify resistance mechanisms that successfully defend against *P. viridiflava* infection. For example, *Marchantia* and other non-flowering plants encode functionally conserved intracellular immune receptors belonging to the nucleotide-binding leucine rich-repeat (NLR) family that often detect type-III effector activity of *P. syringae* pathovars in flowering plants like Arabidopsis [37–39]. Whether the diversity of *P. syringae* and *P. viridiflava* effector alleles are capable of provoking an analogous effector-triggered and NLR-mediated response in liverworts remains to be determined. Looking forward, it is becoming increasingly clear that our understanding both of plant immunity and pathogen virulence will continue to expand beyond the previously limited view that focused only on angiosperm models and crops.

## ACKNOWLEDGEMENTS

We thank the *P. syringae* research community for providing genomic resources that were critical for this work; Jacob Malone (John Innes Centre, UK) for providing the pTS1 vector and protocols for allele replacement in *Pseudomonas*; Connor Forsyth for technical assistance; and all current and past members of the Carella group for insightful discussions on *Marchantia-Pseudomonas* interactions.

## AUTHOR CONTRIBUTIONS

PC designed research; KR, LB, KSC, KG, MH, and PC performed research; MH and PC performed bioinformatic analyses; PC prepared figures and final datasets; KR, MH, WM, and PC supervised research; PC wrote the paper; All authors edited and/or approved the manuscript.

## DATA AVAILABILITY

All gene identifiers and dataset accession numbers are provided in the manuscript. Genomes are deposited at NCBI under the accessions (MPG01, JBITRC000000000; MPG11, JBGMWT000000000; MPG14, JBGMWS000000000; MPG20, JBGMWR000000000; MPG32, JBGMWQ000000000; MPG39, JBGMWP000000000). raw and processed data related to *P. viridiflava* species and virulence factor phylogenies are available online at Zenodo (doi: 10.5281/zenodo.14259570).

## MATERIALS AND METHODS

### Pseudomonas isolation from wild Marchantia

*Pseudomonas* was isolated from natural populations of disease liverworts growing in an urbanized site on the Norwich Research Park campus. Diseased liverwort archegoniophores were harvested in 50 ml conical tubes, surface sterilized with 75% ethanol for 30 seconds with shaking, rinsed with ∼25 ml sterile distilled water 5 successive times, crushed in 3 mL of 10 mM MgSO_4_ buffer, and then serially diluted in 10 mM MgSO_4_ and plated onto *Pseudomonas*-selective KB media supplemented with nitrofurantoin (100 μg ml^-1^). Plates were incubated at 20C for 2 days and 40 single colonies were then streaked onto individual KB-Nitro plates (representing isolates MPG1-40). A single colony from each plate/isolate was grown overnight in KB media and glycerol stocks were prepared and stored at -80 for further use.

### Plant and bacterial growth

The liverwort *Marchantia polymorpha* (Tak-1, Takaragaike-1) was cultivated axenically by placing asexual gemmae clones ontop of sterile nylon mesh (100 micron, Normesh) previously overlaid onto one-half–strength MS-B5 (Murashige and Skoog with B5 vitamins) media (pH 6.7). Plates were sealed with micropore tape and grown under long-day photoperiod (16 hours of light; ∼80 uE light intensity) at 20-22 °C. *Nicotiana benthamiana* were grown in soil under controlled conditions with at 22 °C, 80% relative humidity, and a long day photoperiod (16 hours of light; ∼120-150 μE light intensity). All *Pseudomonas* isolates were grown axenically in King’s B (KB) or low-salt LB (5 g L^-1^ NaCl) media.

### Plant infection assays

*Marchantia polymorpha* Tak-1 plants were infected as described previously[12]. In short, 4-week-old plants grown axenically on nylon mesh overlaid onto ½ MS-B5 agar plates were used for infection assays. Bacteria grown overnight (OD_600_: 0.2-0.8) were harvested by centrifugation, resuspended in 5 mM MgCl_2_ and adjusted to a final concentration of 2×10^6^ colony forming units (CFU) ml^-1^. Bacteria were vacuum infiltrated into Marchantia thalli using negative pressure generated within the chamber of 50 mL syringe. Plants were then placed into petri dishes containing wetted Whatman paper, sealed with micropore tape, and returned to growth conditions until the indicated time points. For *Nicotiana*, fully expanded leaves were pressure infiltrated with inoculum adjusted to 2×10^6^ cfu ml^-1^ and harvested at indicated time points.

### Draft genome sequencing, assembly, and annotation

Draft genome sequencing of *Pseudomonas* isolates derived from *Marchantia* was performed by MicrobesNG (https://microbesng.com). In brief, purified DNA extracted by MicrobesNG was used to generate genomic DNA libraries using the Nextera XT Library Prep kit (Illumina) following the manufacturer’s guidelines. Libraries were subsequently sequenced on an Illumina NovaSeq 6000 using the 250 bp paired end configuraiton. Adapter sequences were trimmed from raw reads using Trimmomatic (v0.30) [40]. De novo assemblies were performed using SPAdes (v03.7) [41] and were annotated using Prokka (v1.11) [42]. Genomes and associated annotations were visualized using Proksee (https://proksee.ca) [43]. Genome information can be found at NCBI (PRJNA1149041) with additional annotation and information available on Zenodo (doi: 10.5281/zenodo.14259570).

### Phylogenetic analyses and virulence factor identification

For phylogenetic placement of MPG isolates into the *P. syringae* species complex, we downloaded the genomes of *P. syringae* type strains [44] alongside a *P. putida* control (Table S1) and performed orthology analyses of proteomes using OrthoFinder2 [45]. Next, we individually aligned each of the 706 orthogroups shared between each bacterial proteome using MAFFT [46] and trimmed the resulting alignments using trimAl [47] under “-strict” mode. The resulting alignments were then concatenated using catfasta2phyml.pl (https://github.com/nylander/catfasta2phyml) and used for maximum likelihood phylogenetic analysis via IQ-TREE[48] v2.0.3 with the “JTT+F+I+G4” model selected through ModelFinder [49] and with 1000 ultrafast bootstrap replicates [50]. The phylogenetic tree was subsequently rendered using TVbot [51].

The *pel* gene was identified/annotated directly by Prokka. By comparison, Type-III effectors were not annotated and only identified through tblastn and/or BLASTP [52] analysis of an exhaustive set of *P. syringae* effector sequences [53] against each MPG genome. Candidate PEL homologs were identified from the UniProt reference database by searching for proteins with at least 50% sequence similarity to *P. viridiflava* PEL (Q4JRB8). For each identified virulence factor, proteins were aligned with MAFFT, trimmed with trimAl, and phylogenies were built using IQ-TREE [48] with ModelFinder [49] and 1000 ultrafast bootstraps [50]. The models selected by ModelFinder were: “JTT +G4” (AvrE, HopAA1, HopM1), “JTT+I+G4” (HopA1), “JTT+F+G4” (HopB2), or “LG+G4” (PEL). Phylogenetic trees were rendered in iTOL or TVBot [51].

### *Pseudomonas viridiflava* allele replacement

Allele replacement was performed in the MPG32 isolate background using the suicide vector pTS1 as previously described [54]. In brief, 1000 bp fragments encoding the 500 bp upstream and 500 bp downstream sequences flanking the genes of interest were synthesized and cloned directly into pTS1 by Twist Bioscience. Constructs (Table S2) were electroporated into MPG32 following an established *Pseudomonas* protocol [55]. Single crossover integrations were first selected on L media containing tetracycline, were re-streaked onto selective media, and then grown overnight in L media without selection. Counterselection to identify double crossovers was performed by serial dilution plating on L media supplemented with 10% sucrose. Sucrose insensitive colonies were tested for tetracycline (12.5 µg ml^-1^) susceptibility (loss of integration) and deletions were confirmed by PCR using the primers listed in Table S3.

### Pectate lyase activity assay

Pectate lyase activity was inferred by the production of soft-rotting when isolates were co-cultivated on potato slices [56]. In brief, store bought potatoes were briefly washed in water and then submerged in 70% ethanol for 10 minutes. Potatoes were then washed at least 5 times with sterile double distilled water before being sliced into approximately 1 cm thick slices. Each slice was individually placed into a sterile petri dish containing 3 layers of wetted Whatman paper. Bacteria were inoculated onto the slice by dipping sterile toothpicks into freshly grown colonies and pricking an x-shaped pattern into the center of the slice. Plates were then sealed with parafilm and kept at room temperature on lab benches for 3 days before imaging.

## SUPPLEMETARY INFORMATION

**Figure S1. *Marchantia* reinfection assays against wild-isolated *Pseudomonas* MPG strains**

**Figure S2. >MPG draft genome characteristics**

**Figure S3. AvrE phylogeny**

**Figure S4. HopM1 phylogeny**

**Figure S5. HopAA1 phylogeny**

**Figure S6. HopA1 phylogeny**

**Figure S7. HopB2 phylogeny**

**Figure S8. PEL phylogeny**

**Figure S9. PCR validation of *P. viridiflava* MPG knockout strains**

**Table S1. Genome resources used in this study**

**Table S2. Vectors used in this study**

**Table S3. Primers used in this study**

