## Supplementary Figures, Tables, and Datasets for "Conserved effectors underpin the virulence of liverwort-isolated *Pseudomonas* in divergent plants": SupportingFigures.pdf

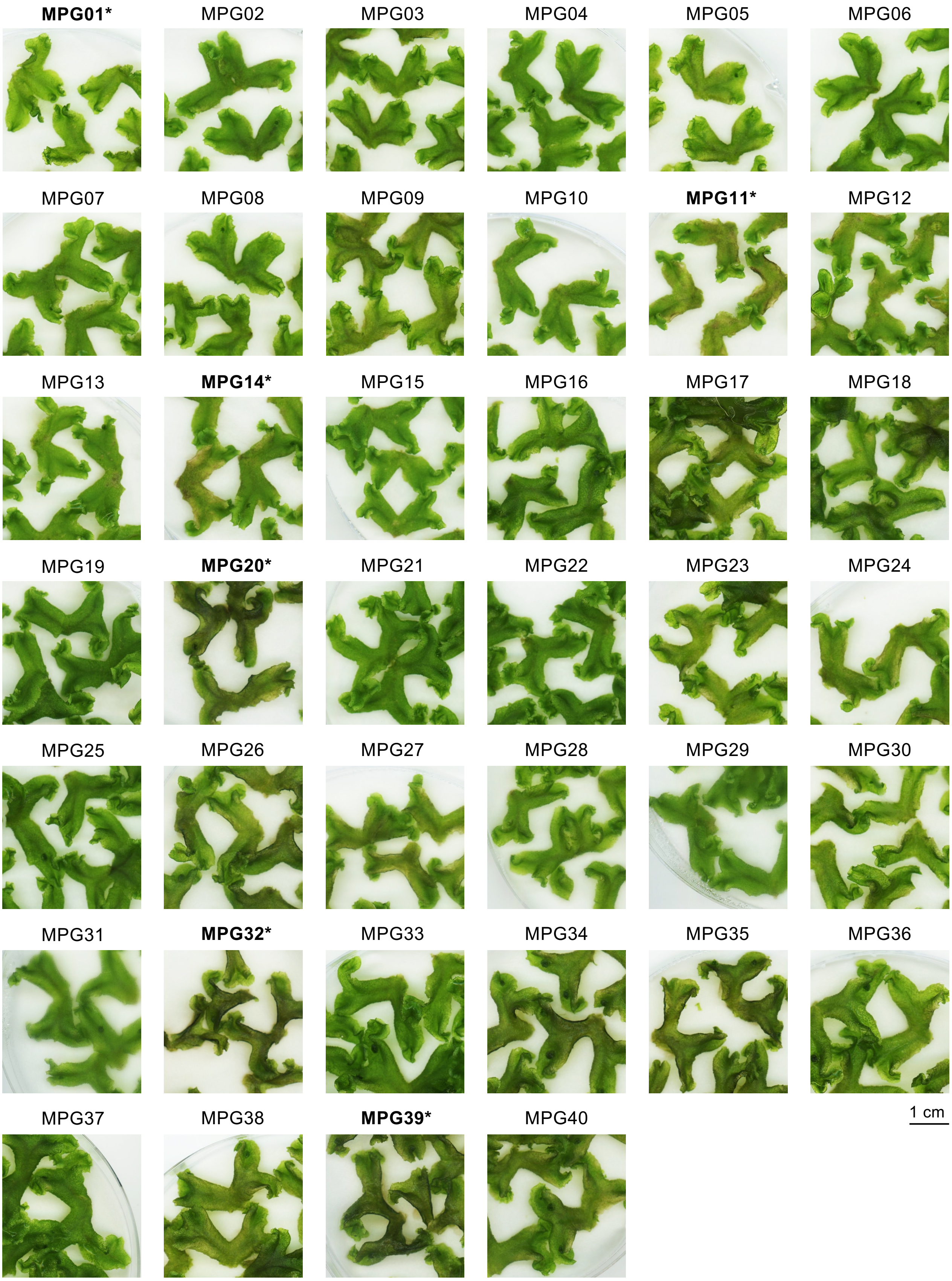


**Figure S1. Marchantia reinfection assays against wild-isolated *Pseudomonas* MPG strains**
Symptom development of axenically grown wildtype *Marchantia polymorpha* (Tak-1; 4-week-old thalli) infected with 40 individual MPG isolates. Symptoms were recorded 7 days post infection (dpi) (n ≥ 8). Isolates used in further experiments are indicated by an asterisk and bold text. Scale bars = 1 cm.

**A**

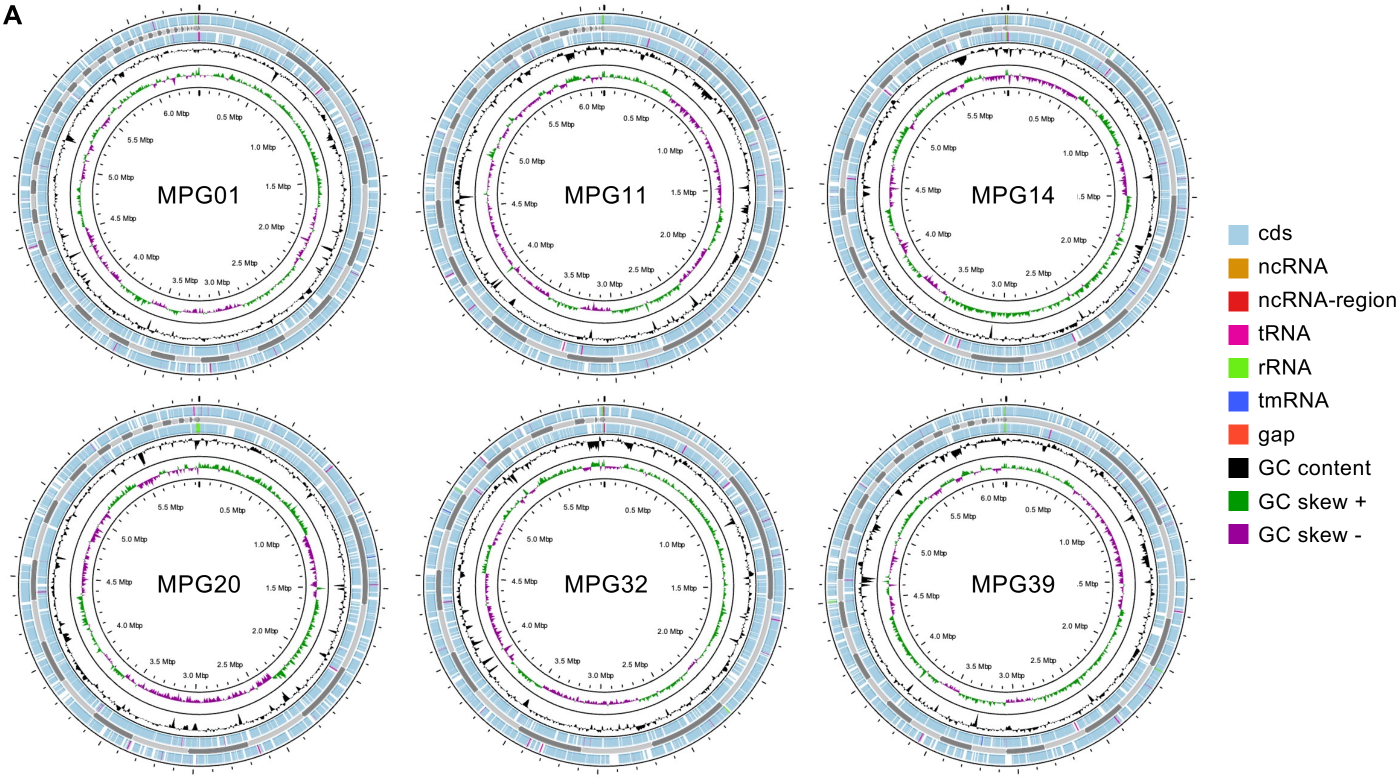


**B**

| Isolate | Sequencing Identifier | Genome size (bp) | Contigs | Features |
|---|---|---|---|---|
| MPG01 | 248141E | 6,399,976 | 96 | 5686 |
| MPG11 | 248142E | 6,124,258 | 65 | 5490 |
| MPG14 | 248143E | 5,931,614 | 67 | 5336 |
| MPG20 | 232580E | 5,936,815 | 74 | 5328 |
| MPG32 | 232581E | 5,931,516 | 60 | 5367 |
| MPG39 | 248144E | 6,126,881 | 72 | 5493 |

**Figure S2. MPG draft genome characteristics**

(A) Draft genome maps of MPG01, MPG11, MPG14, MPG20, MPG32, and MPG39. Common genomic features are displayed for each strain.

(B) Overview of MPG isolate genome characteristics. Genomic features were annotated using Prokka (as described in the methods).

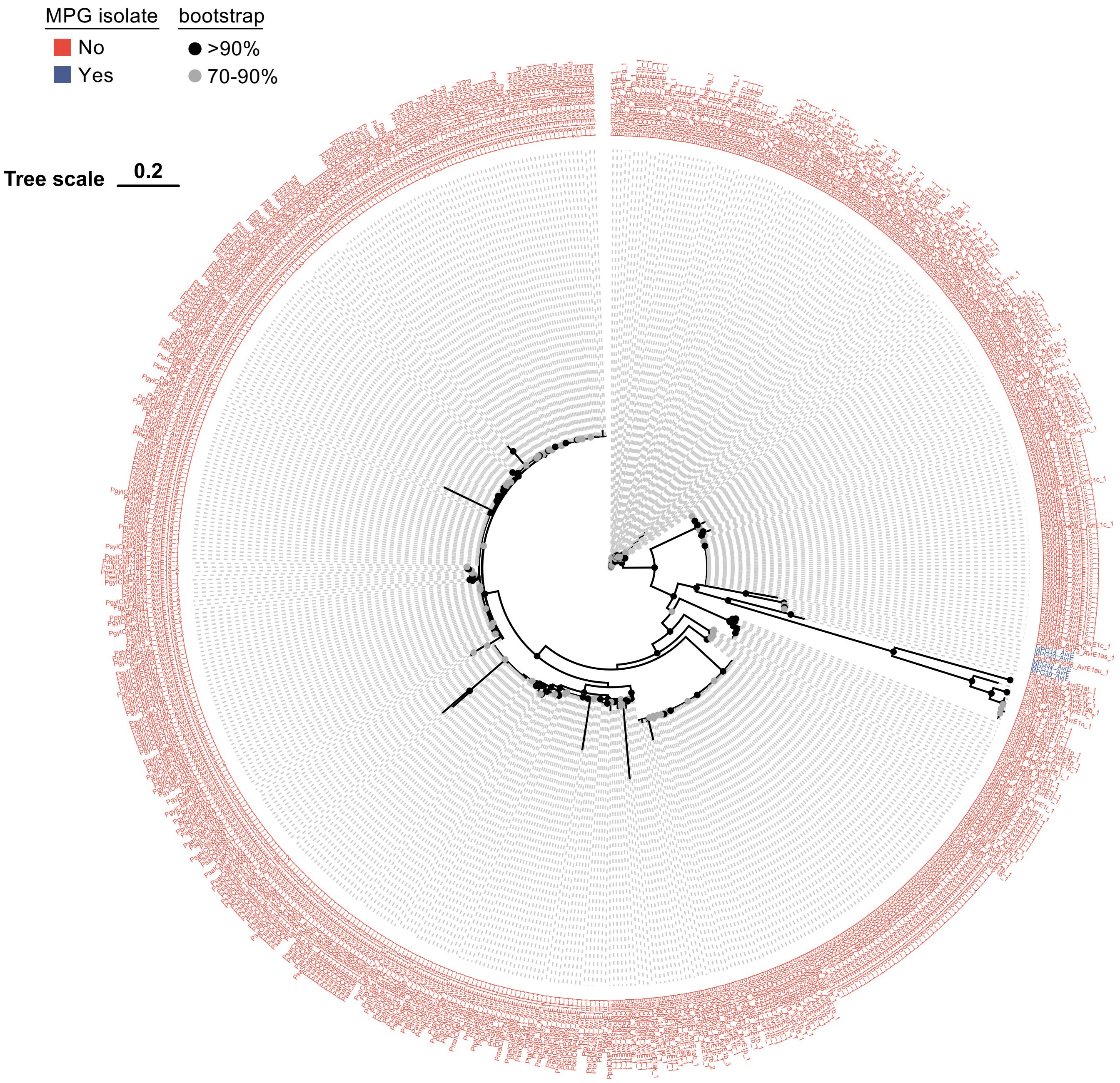


**Figure S3. AvrE phylogeny**

Maximum likelihood phylogeny of AvrE protein sequences from MPG isolates (blue) compared against those from diverse *Pseudomonas syringae* species complex members (red) with bootstrap confidence value ranges indicated. Tree scale = substitutions/site.

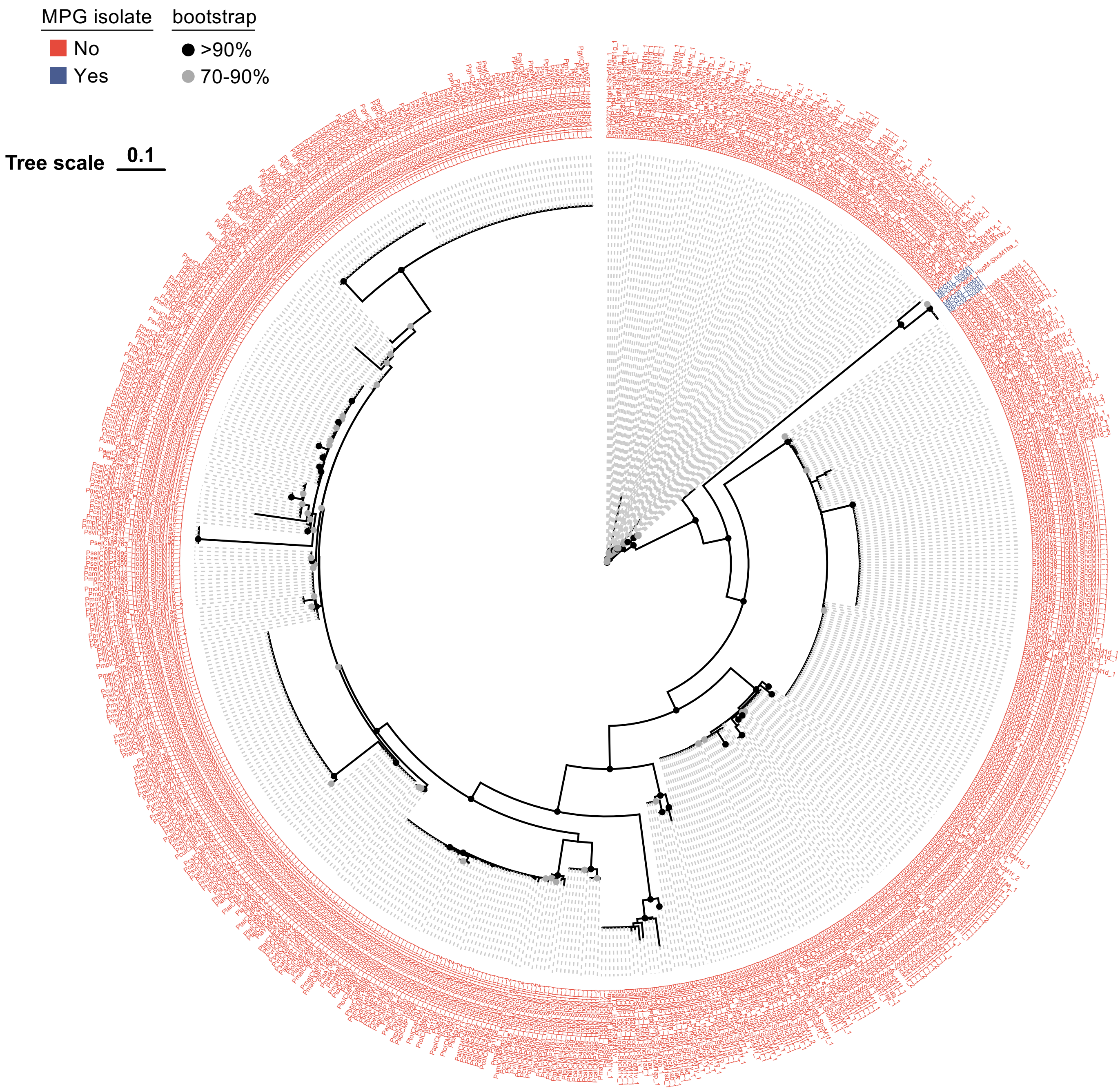


**Figure S4. HopM1 phylogeny**

Maximum likelihood phylogeny of HopM1 protein sequences from MPG isolates (blue) compared against those from diverse *Pseudomonas syringae* species complex members (red) with bootstrap confidence value ranges indicated. Tree scale = substitutions/site.

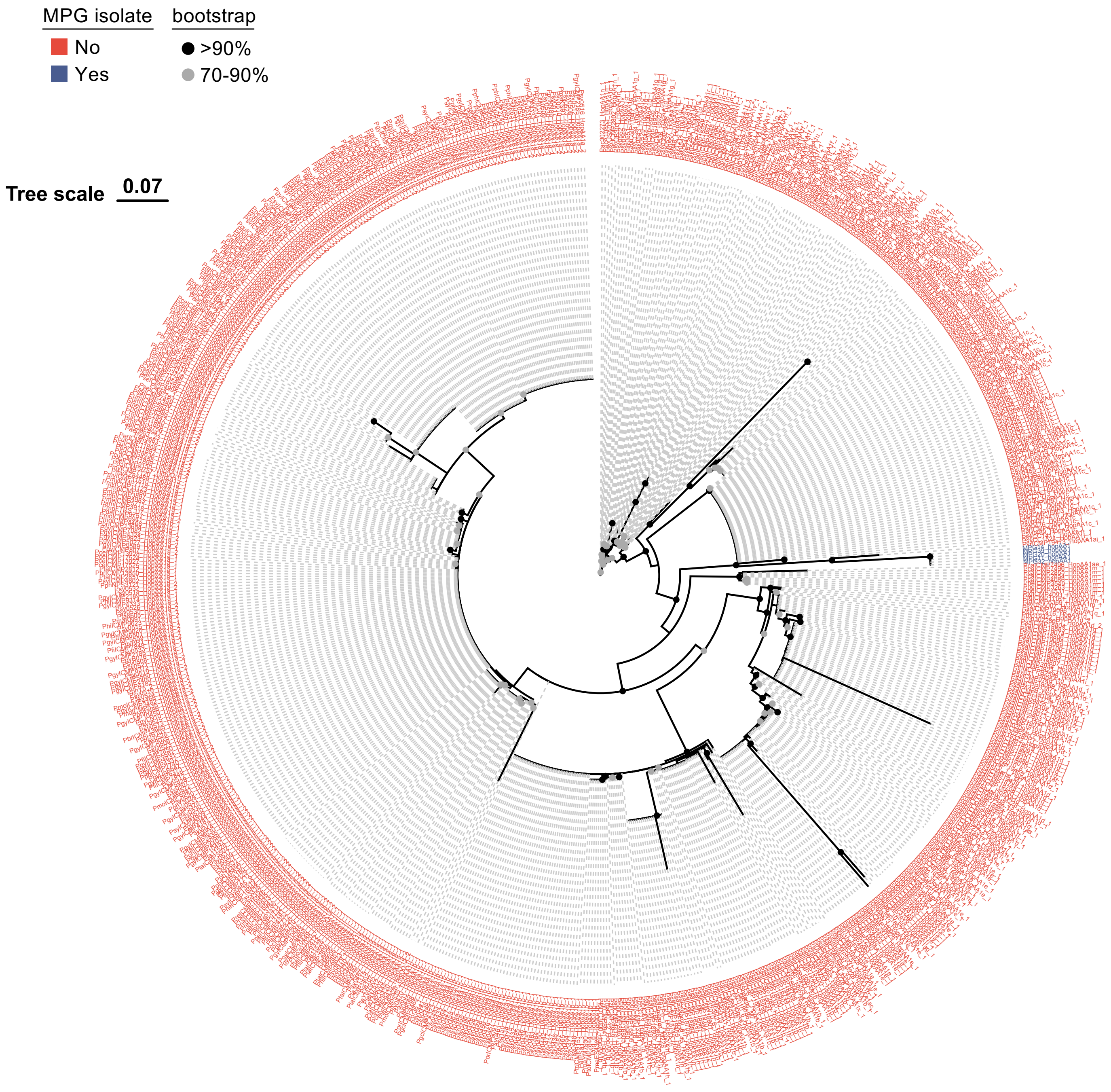


**Figure S5. HopAA1 phylogeny**

Maximum likelihood phylogeny of HopAA1 protein sequences from MPG isolates (blue) compared against those from diverse *Pseudomonas syringae* species complex members (red) with bootstrap confidence value ranges indicated. Tree scale = substitutions/site.

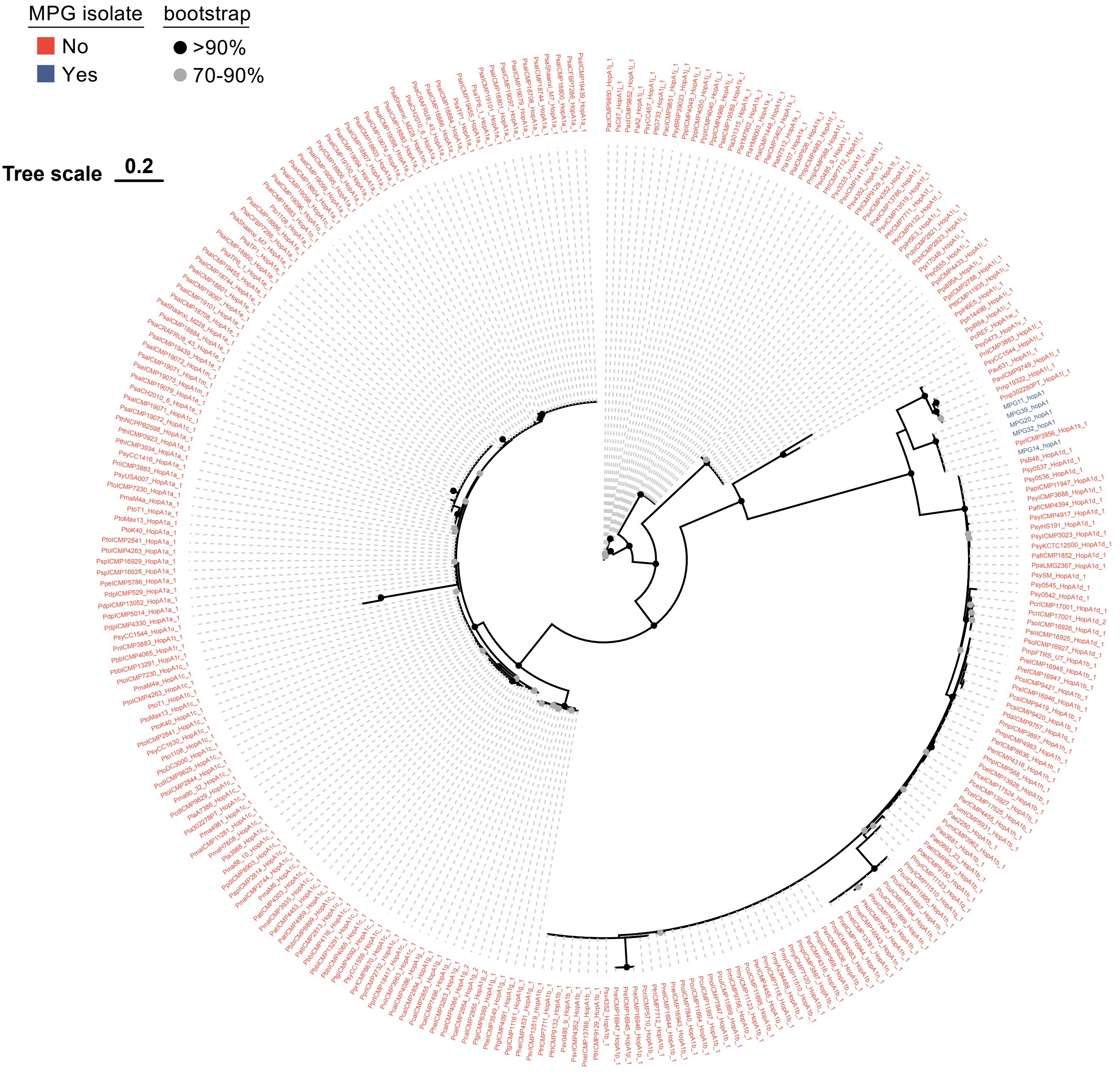


**Figure S6. HopA1 phylogeny**

Maximum likelihood phylogeny of HopA1 protein sequences from MPG isolates (blue) compared against those from diverse *Pseudomonas syringae* species complex members (red) with bootstrap confidence value ranges indicated. Tree scale = substitutions/site.

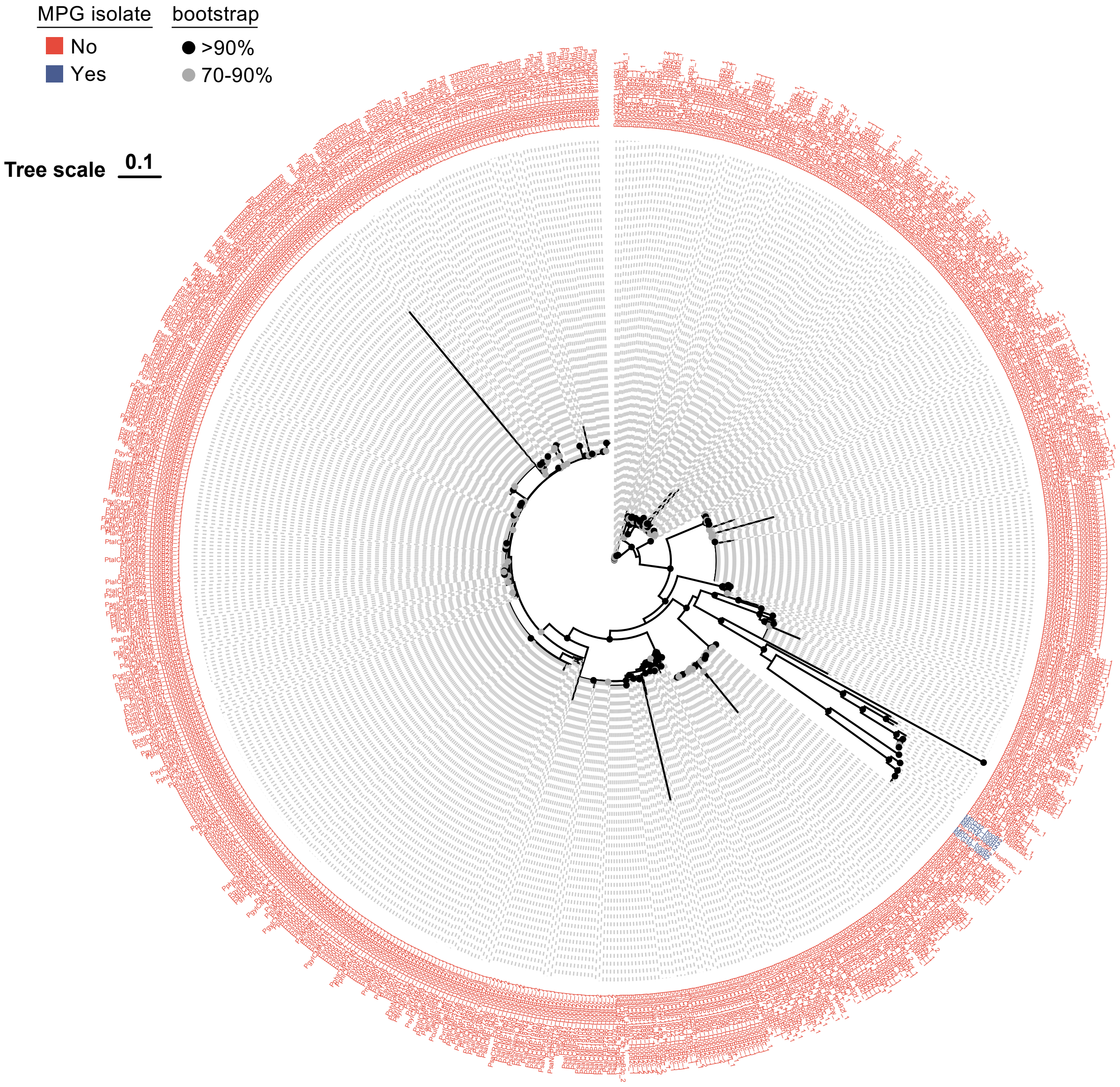


**Figure S7. HopB2 phylogeny**

Maximum likelihood phylogeny of HopB2 protein sequences from MPG isolates (blue) compared against those from diverse *Pseudomonas syringae* species complex members (red) with bootstrap confidence value ranges indicated. Tree scale = substitutions/site.

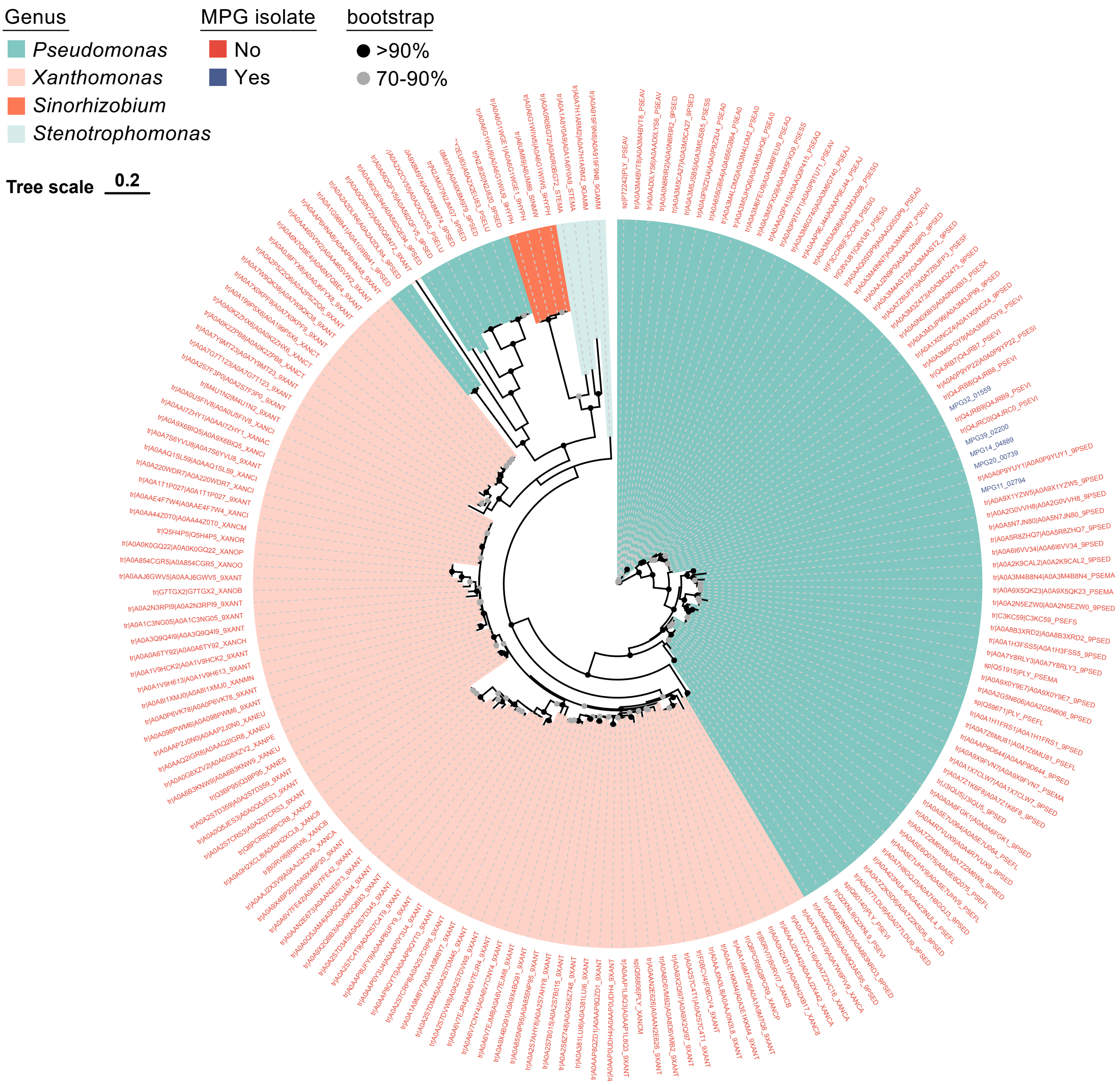


**Figure S8. PEL phylogeny**

Maximum likelihood phylogeny of PEL protein sequences from MPG isolates (blue) compared against those from diverse bacteria (Pseudomonas, Xanthomonas, Sinorhizobium, Stenotrophomonas) with bootstrap confidence value ranges indicated. Tree scale = substitutions/site.

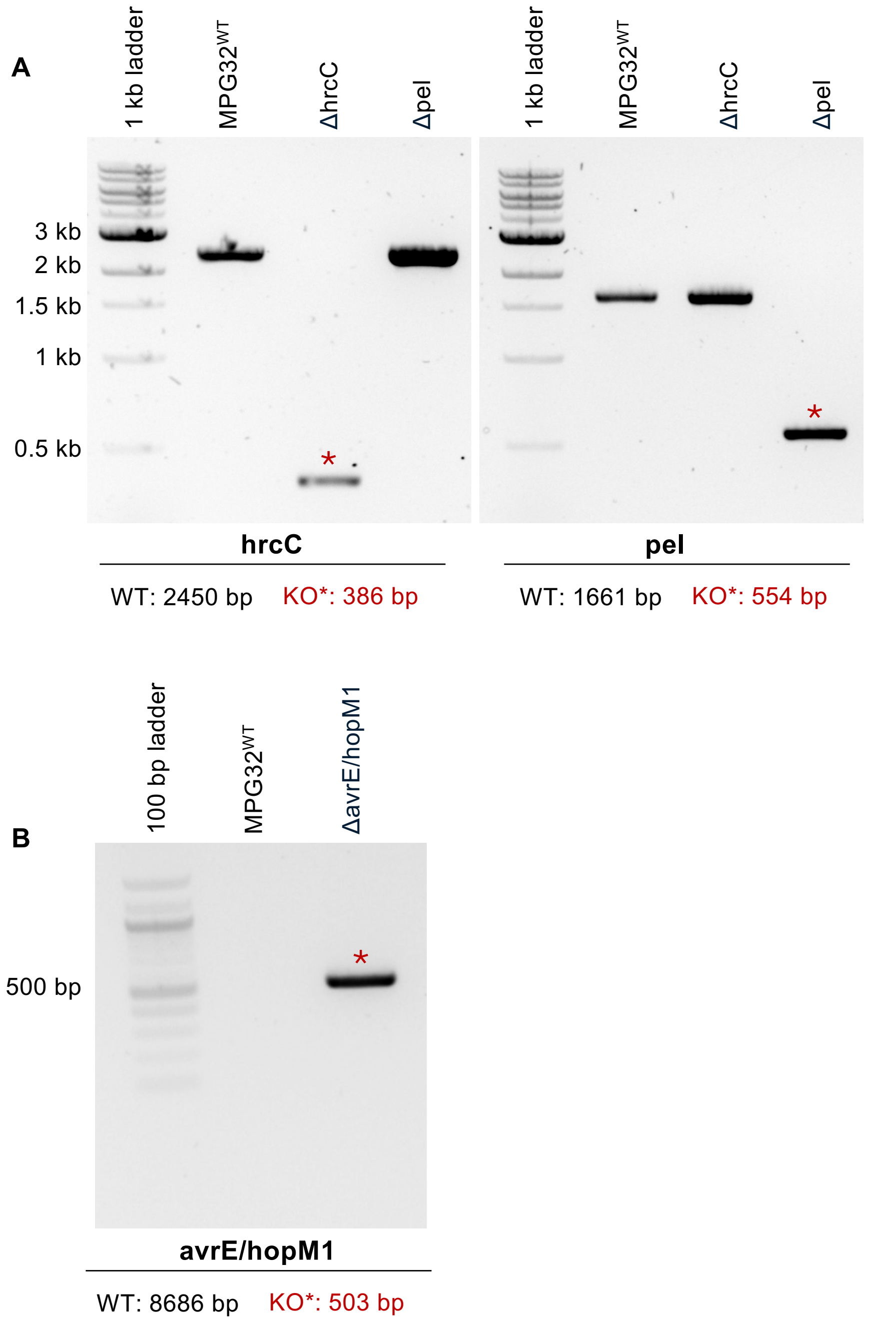


**Figure S9. PCR validation of *P. viridiflava* MPG knockout strains**

(A) PCR validation of hrcC or pel gene knockouts in *Pseudomonas viridiflava* MPG32 isolates. The different sizes of PCR products produced by wild-type MPG32 (WT) or each respective knockout (KO) mutant (ΔhrcC or Δpel) are indicated below each gel image. Amplicons representative of successful gene knockouts are indicated by an asterisk *.

(B) PCR validation of the ΔavrE/hopM1 knockout in *Pseudomonas viridiflava* MPG32. The different sizes of PCR products produced by wild-type MPG32$^{WT}$ or ΔavrE/hopM1 are indicated below the gel image. The amplicon representing successful knockout is indicated by an asterisk *.
